# Cryo-EM structures of human α1B/βI+βIVb microtubules shed light on isoform specific assembly

**DOI:** 10.1101/2023.12.01.569594

**Authors:** Elena A. Zehr, Antonina Roll-Mecak

## Abstract

Microtubules are dynamic polymers assembled from αβ-tubulin dimers. Mammals have multiple α and β-tubulin isoforms. Despite a high degree of conservation, microtubules assembled from different tubulin isoforms have unique dynamic properties. How isoform sequence variation affects polymerization interfaces and nucleotide dependent conformational changes in microtubules is still not well understood. Here we report 2.9Å resolution cryo-EM structures of human α1B/βI+βIVb microtubules in the GDP state, and a GTP-like state, bound to GMPCPP. Our structures show that, similar to microtubules assembled from other mammalian isoforms, transition from the GTP to the GDP states in α1B/βI+βIVb microtubules results in strengthening of the longitudinal interface and an overall compaction of the axial dimer repeat distance in the lattice. Interestingly, we find that α-tail residues link longitudinally adjacent tubulin dimers through interactions with two conserved arginine residues in β-tubulin that are mutated in human disease. Comparative analysis of tubulin isoforms shows minimal isoform-specific effects at the longitudinal interface or the α-tubulin lateral interface, but a high concentration of sequence variability in the second shell of residues away from the β-tubulin lateral interface which can modulate polymerization interfaces and thus impact microtubule dynamics.

## Introduction

Microtubules are essential for eukaryotic cell motility, cell division and intracellular transport. Microtubules consist of αβ-tubulin dimers that assemble longitudinally in head-to-tail fashion into polar protofilaments that then associate laterally to form the cylindrical microtubule. Microtubules are dynamic; they undergo stochastic phases of growth and shrinkage at their ends termed “dynamic instability”^1,2^. This dynamic behavior is driven by the GTPase cycle of the αβ-tubulin dimer. The tubulin dimer binds two GTP molecules. β-tubulin at the microtubule plus end contains the GTP exchangeable site, known as the E-site. The GTP binding site on β-tubulin is exposed and GTP is hydrolyzed at this site during polymerization. α-tubulin at the minus end contains the non-exchangeable and non-hydrolysable GTP site. This site is permanently bound to GTP during polymerization and depolymerization. Cryo-EM structures of microtubules bound to GDP and to the non-hydrolysable analogs GMPCPP and GTPγS ^3–5^ revealed how GTP hydrolysis and phosphate release in the β-tubulin E site remodels the active site and leads to conformational changes at the longitudinal interdimer interface (the interface between neighboring dimers in a protofilament) that results in an overall compaction of the microtubule dimer repeat ^3–6^. However, the compaction of the microtubule lattice between the GTP and GDP bound states is not universal. For example, cryo-EM structures of *S. cerevisiae* microtubules revealed that they do not undergo lattice compaction ^7^.

The tubulin dimer is highly conserved in all species, consistent with its indispensable role in the survival of eukaryotes and the fact that most of the protein is involved in either GTP binding or polymerization interfaces. However, organisms express multiple tubulin isoforms ^8,9^ that have non-overlapping functions ^10^. Different structures and dynamic parameters have now been reported even for the highly similar human tubulin isoforms^11,12–15^ indicating that subtle sequence differences can result in functional differences. Tubulin isoforms display the highest sequence variation in two functionally important regions: the interface between neighboring protofilaments, also known as the lateral interface, and the intrinsically disordered C-terminal tails ^9^. The sequence and conformational variability at the lateral interfaces were proposed to be responsible for isoform specific dynamic properties and distinct microtubule lattice architectures ^11–13,16,17^. The C-terminal tubulin tails perform regulatory functions and interact with many motors and microtubule associated proteins ^10^. They are disordered in all cryo-EM structures of undecorated microtubules to date and have been visualized only when the microtubule is bound to an effector such as a tubulin modifying enzyme ^18^ or the NDC80 complex ^19^.

While Cryo-EM studies in the last three decades have provided fundamental insights into how tubulin assembles into microtubules and the GTP hydrolysis driven conformational changes in the microtubule lattice that underlie dynamic instability ^3–5^, our understanding about isoform specific lattice interactions is still in its infancy. Notably, many of the polymerization interfaces display sequence variability among tubulin isoforms and thus subtle changes at these interfaces could impact the energetics of lattice assembly and disassembly. Here we report 2.9Å resolution structures of α1B/βI+βIVb microtubules assembled from endogenous tubulin purified from human embryonic kidney (tsA201) cells in two nucleotide states. Comparison of structures in a GTP-like and GDP state show, that similar to undecorated brain microtubules^20^ and microtubules assembled from α1A/βIII recombinant tubulin ^6^, α1B/βI+βIVb microtubules also undergo conformational changes upon GTP hydrolysis and phosphate release which strengthen the longitudinal interface and result in an overall compaction of the microtubule lattice. Thus, lattice compaction seems to be a broadly shared GTP hydrolysis responsive structural rearrangement in mammalian microtubules ^3–5,6,20^. Interestingly, our high-resolution reconstruction revealed additional density for the N-terminal section of the α-tail and shows that residues in the α-tail link longitudinally adjacent tubulin dimers through interactions with two conserved arginine residues in β-tubulin which are mutated in human disease. Detailed analysis of the β-tubulin lateral interface, a hotspot for sequence variability, sheds light on how isoform sequence variations can modulate polymerization interfaces and thus impact microtubule dynamics.

## Results

### Lattice compaction between GMCPPP and GDP bound states

We purified tubulin from the epithelial-like transformed embryonic kidney cell line tsA201 using a TOG1 affinity purification as previously described ^21^. This tubulin preparation contains predominantly one α-tubulin isoform, α1B, and two β-tubulin isoforms, βI and βIVb ^12^, a composition characteristic of fibroblasts and other commonly used laboratory cell lines ^22^. To gain insight into isoform specific contacts and GTP-hydrolysis induced structural changes, we solved cryo-EM structures of α1B/βI+βIVb microtubules bound to GDP and the GTP-analogue guanylyl-(α,β)-methylene-diphosphonate (GMPCPP) to 2.9 Å (Figs. 1(a), (b), Fig. S1 and Fig. S2). Similar to brain microtubules ^20^, the majority of α1B/βI+βIVb microtubules are 14-stranded: ∼55% and ∼80% for the GDP and GMPCPP, respectively (Fig. S1 (b)). Since microtubule lattices incorporate distortions that can limit the resolution of cryo-EM reconstruction, we used a protofilament refinement procedure which takes into account lattice flexibility ^23^ (Methods). This is the highest resolution achieved so far for a non-brain microtubule structure and provides detailed information into isoform specific microtubule assembly. Our previous structure of α1B/βI+βIVb microtubules was resolved only to 4.2 Å ^12^, limiting analysis of side-chain contacts at polymerization interfaces. The nucleotides in both α- and β-tubulin are well-defined (Figs. 1(c-f)). In the GMPCPP assembled microtubules (Methods), the GMPCPP in the E-site is well-resolved, with all three phosphates clearly defined as well as the bound Mg^2+^ ion (Fig. 1(e)). Likewise, the structure of a microtubule assembled in the presence of GTP, shows the GDP in the E-site, with both the α- and β-phosphate clearly defined (Fig. 1(f)). The axial dimer repeat distance in α1B/βI+βIVb microtubules compacts between the GMPCPP and GTP-bound states, changing from 84.22 ± 0.08 Å to 81.67 ± 0 Å (p < 0.0001 by t-test; Fig. 1 (g), Fig. S2(c)). Thus, lattice compaction seems to be a broadly shared GTP hydrolysis responsive structural rearrangement in mammalian microtubules, observed for porcine brain microtubules ^3–5^, recombinant human a1A/ýIII microtubules ^6^ and now human α1B/βI+βIVb microtubules.

**Figure 1.**
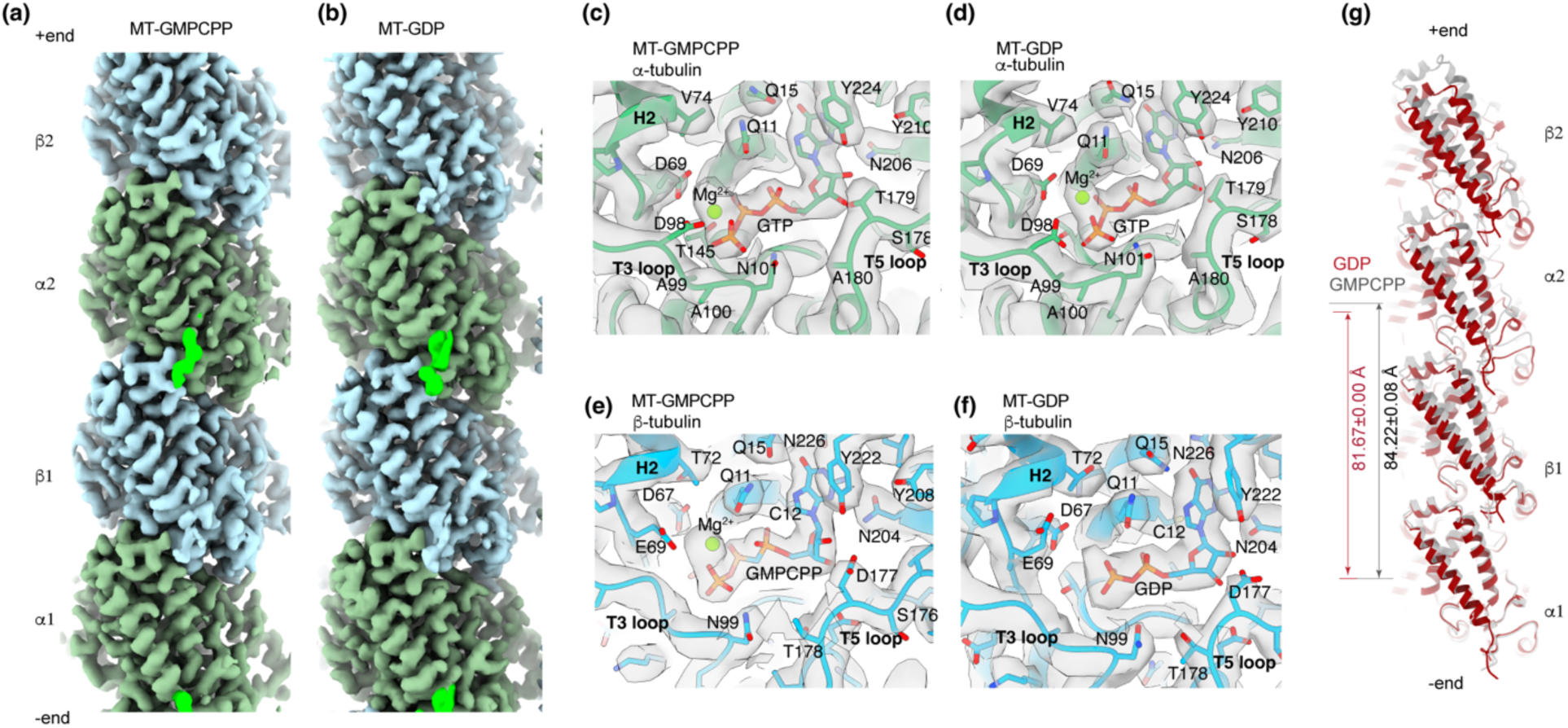
Cryo-EM reconstructions of human α1B/βI+βIVb microtubules bound to GDP and GMPCPP. **(a-b)**. Cryo-EM maps of 2 tubulin dimers within a protofilament from GMPCPP-(a) and GDP-bound (b) α1B/βI+βIVb microtubules, respectively; ý-tubulin, blue and a-tubulin, green. Microtubule minus and plus ends indicated. The density corresponding to the N-terminus of the a-tubulin tail is in bright green. **(c-d)** Views of the a-tubulin nucleotide-binding site in GMPCPP-(c) and GDP-microtubules (d), respectively. **(e-f)**. Views of the ý-tubulin nucleotide-binding site in GMPCPP-(e) and GDP-microtubules (f), respectively. **(g).** Single protofilament of GMPCPP- and GDP-microtubules aligned on a1-tubulin showing compaction of the lattice in response to hydrolysis and phosphate release with the axial dimer repeat changing from 84.22±0.08 Å in GMPCPP-to 81.67±0 Å in GDP-microtubules (Methods).

### a-tails link tubulin dimers along the protofilament

Our high-resolution reconstructions revealed clear additional density for α-tail residues 439-441 (Figs. 1(a-b)). Interestingly, this region of the α-tubulin tails interacts in-trans with the body of the β-tubulin protomer of the longitudinally adjacent tubulin dimer (Fig. 2(a-c)). Specifically, R390 in H11 of β1 is within H-bonding distance to the backbone of V437 and S439 in the tail of α2 (the longitudinally adjacent dimer), while R391 in H11 of β1 interacts with E441 in the α2-tail (Fig. 2(a-b)). The contacts with the backbone of the α-tail would ensure interactions with α-tail sequences from different isoforms ^24^ (Fig. S3). These contacts between the α-tails with the longitudinally adjacent tubulin dimer at the minus end augment the “canonical” longitudinal interface stabilizing the microtubule. No density for the α-tail is visible beyond residue 442, consistent with a high conformational heterogeneity beyond this point. The trans interaction of the α-tail with the longitudinally adjacent dimer is also visible in the GDP-bound structure, however the density for the α-tail is slightly less defined (Fig. 2(c)), most likely due to the smaller number of particles used in this reconstruction (Fig. S1(c)). Consistent with their importance at this interface, both R390 and R391 in β-tubulin are invariant among mammalian tubulin isoforms (Fig. S4) and also mutated in human disease. R390 is mutated to glutamine in patients with Uner Tan syndrome ^25^, a condition characterized by quadrupedalism, mental retardation and cerebellar defects. R391 is mutated to histidine or serine in patients with Leber congenita amaurosis ^26^, a condition characterized by severe visual impairment at birth.

**Figure 2.**
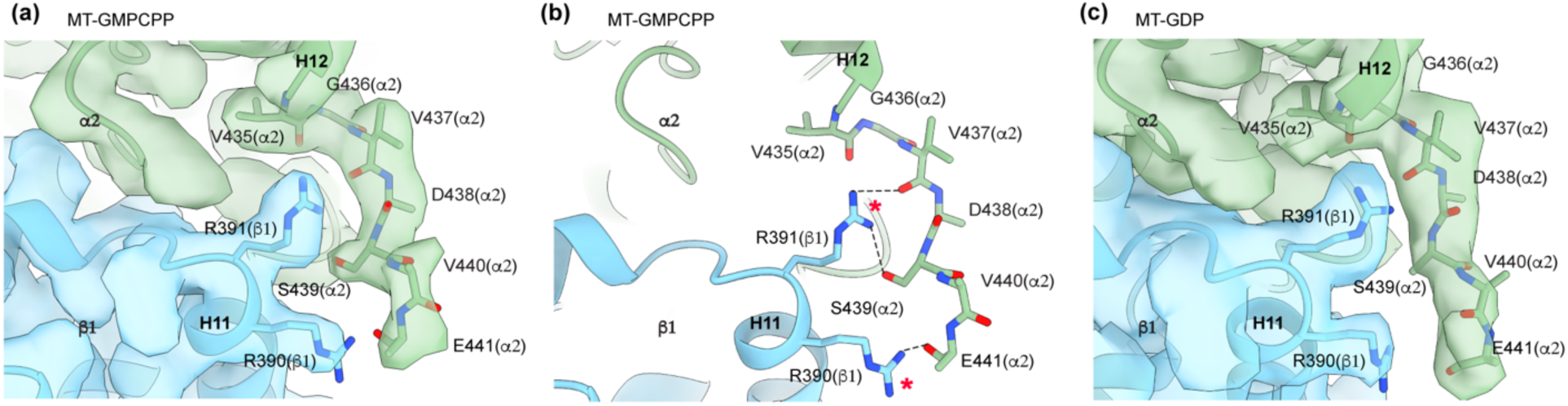
a-tail residues interact with the β-tubulin body of the longitudinally adjacent dimer. **(a-b)**. Views showing the interaction between the α2−tubulin C-terminal tail and the β1-tubulin body in GMPCPP microtubules. Cryo-EM map (a), transparent, colored green and blue for α-and β-tubulin, respectively. α2-tubulin tail residues, shown as sticks and color-coded by heteroatom, interact with β1-tubulin residues R390 and R391 (b). Hydrogen bonds shown as black dashed lines. Residues mutated in human disease highlighted by red stars^44^. **(c)**. View showing the interaction between the α2−tubulin C-terminal tail and the β1-tubulin body in a GDP-microtubules. Cryo-EM map and atomic model colored and represented as in (a).

### Lattice compaction and increase in the longitudinal interface between the GMPCPP and GDP states

Each tubulin protomer contains a N-terminal nucleotide-binding domain (NBD), proximal to the microtubule plus end, an intermediate domain (ID) proximal to the minus end and connected to the NBD through helix H7, and a C-terminal domain (CTD) that lines the microtubule outer surface and also contains the disordered C-terminal tails ^27^. The GTP in a-tubulin is bound at the aý interdimer interface and the GTP in ý-tubulin is bound at the aý intradimer interface. Hydrolysis of the GTP at the E site in ý-tubulin is catalyzed by the invariant E254 of H8 supplied in trans by a-tubulin when a new tubulin dimer is added at the plus end of the protofilament ^27,28^. To assess E site nucleotide-induced structural rearrangements, we aligned the β1-tubulin protomers in the GDP and GMPCPP-bound states using the rigid structural elements that coordinate the nucleotide base (residues 10-20 and 221-227) (Fig. 3(a)). This showed that loss of the γ-phosphate at the E site pulls the T7 loop in α2-tubulin and helix H8 that contains the catalytic E254 ∼ 1.5 Å closer to β1-tubulin (Fig. 3(b-c)). These structural rearrangements proximal to the nucleotide binding pocket induce an overall downward movement of the entire α2 protomer by ∼2.6 Å. The nucleotide-coordinating T3 and T5 loops in β1-tubulin also move closer to this interface due to the loss of the γ-phosphate (Fig 3(d)). Overall, these nucleotide-driven conformational changes result in a compaction at the longitudinal interface between β1 and α2. To assess the effects of these nucleotide-induced structural rearrangements on α-tubulin, we aligned the α2-tubulin protomers in the GDP and GMPCPP bound microtubule structures using the rigid structural elements that coordinate the nucleotide base. This alignment shows that in response to hydrolysis and phosphate release at the E site, the NBD and ID+CTD domains in α2 rotate 3°, with the latter two domains rotating in a concerted motion outwards from the lumen towards the microtubule surface (Fig. 2(e)). Similar structural rearrangements have been described for brain microtubules ^3,5^, and thus are shared across multiple mammalian tubulin isoforms in response to the nucleotide state.

**Figure 3.**
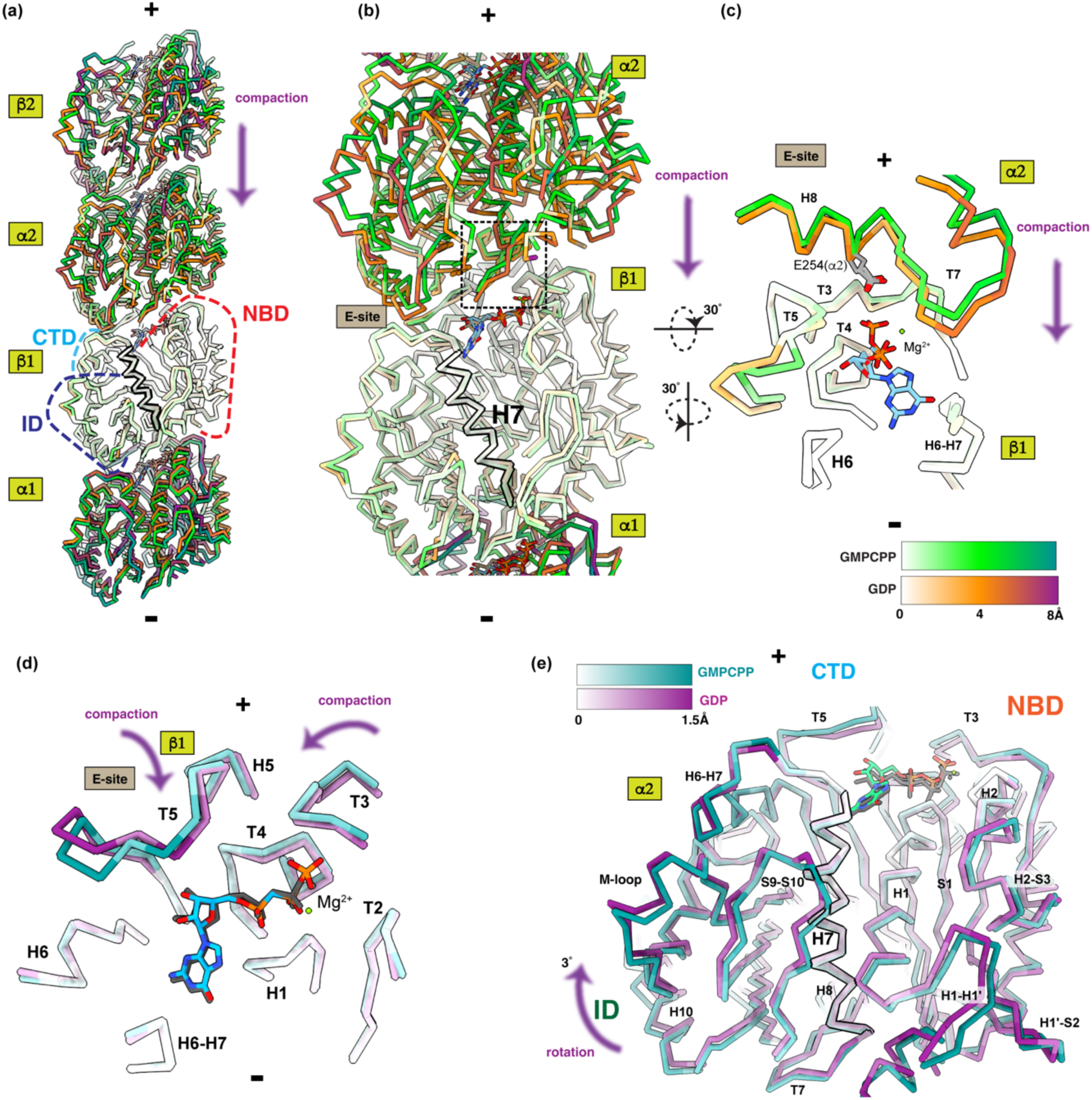
Nucleotide-induced structural rearrangements in the E-site propagate along the protofilament. **(a)** GDP and GMPCPP-bound protofilament models, shown as chain trace, are aligned on structural elements coordinating the nucleotide base bound to β1-tubulin. Models colored by RMSD between αCs; GMPCPP structure colored on a gradient from white to bright green to dark green; GDP structure colored on a gradient from white to orange to maroon. “+” and “-“indicate the plus and minus ends, respectively. Nucleotide binding domain, NBD, outlined with a red dashed line; intermediate domain, ID, dark blue dashed line; central helix H7, black outline; C-terminal domain, CTD, light blue dashed line. (**b**) The E-site with the catalytic T7 loop and helix H8 highlighted with a black dashed line. (**c**) Lattice compaction brings the catalytic E254 residue in H8 of α2 closer to the β1-tubulin nucleotide binding site. (**d**) Movement of the β1 nucleotide-coordinating loops between the GMPCPP and GDP-bound states. **(e)**. 3° rotation of ID and the H1-H1’ loop in the NBD in the α2-tubulin protomer. α2-tubulin subunits from GDP and GMPCPP-bound protofilament models were aligned to each other using structural elements that coordinate the nucleotide base. The models are colored by RMSD between αCs; GMPCPP structure colored on a gradient from white to dark cyan, GDP structure colored on a gradient from white to maroon.

Transition to the GDP state also leads to a significant increase in the longitudinal interface. The total area buried at the longitudinal interface changes from 1422.3 Å^2^ in the GMPCPP-bound state to 1803.6 Å^2^ in the GDP state (Fig. 4(a)). In the GDP state, the S2-H2, T5 and H6-H7 loops in β1-tubulin, involved in nucleotide binding, and the H11-H12 loop, make three novel contact points with α2-tubulin, compared to the GMPCPP structure (Fig. 4(a-c)). The first set of contacts is made between E69 in the S2-H2 loop of β1 with D251 in α2, and between and P70 and M1 and R2 in α2-tubulin (Fig. 4(a-c), Fig. S4, Fig. S4). The second contact is between K174, V180 and P182 in the T5 loop of β1 with A333 in helix H10, T257 in H8 and T349 in S9, respectively (Fig. 4(a-c), Fig. S3, Fig. S4). The third new contact is between F212 in H6, T219 and T221, both in the H6-H7 loop of β1 with K326 in H10, and V324 and P325 in the S8-H10 loop of α2 tubulin (Fig. 4(a-c), Fig. S3, Fig. S4). Except for F212 which is substituted to serine in βVIII (TUBB8), all the residues that make differential contacts between the GMPCPP and GDP-bound states are invariant in all human tubulin isoforms indicating a very high penalty for any substitutions at this interface (Fig. S3, S4). The F212 substitution to serine in βVIII (TUBB8) would weaken the interface in the GDP-bound state. Interestingly, F212 interacts with K326, which is mutated to asparagine in patients with microlissencephaly^29^. Many of the residues at the longitudinal interface are mutated in human disease (Fig. 4b).

**Figure 4.**
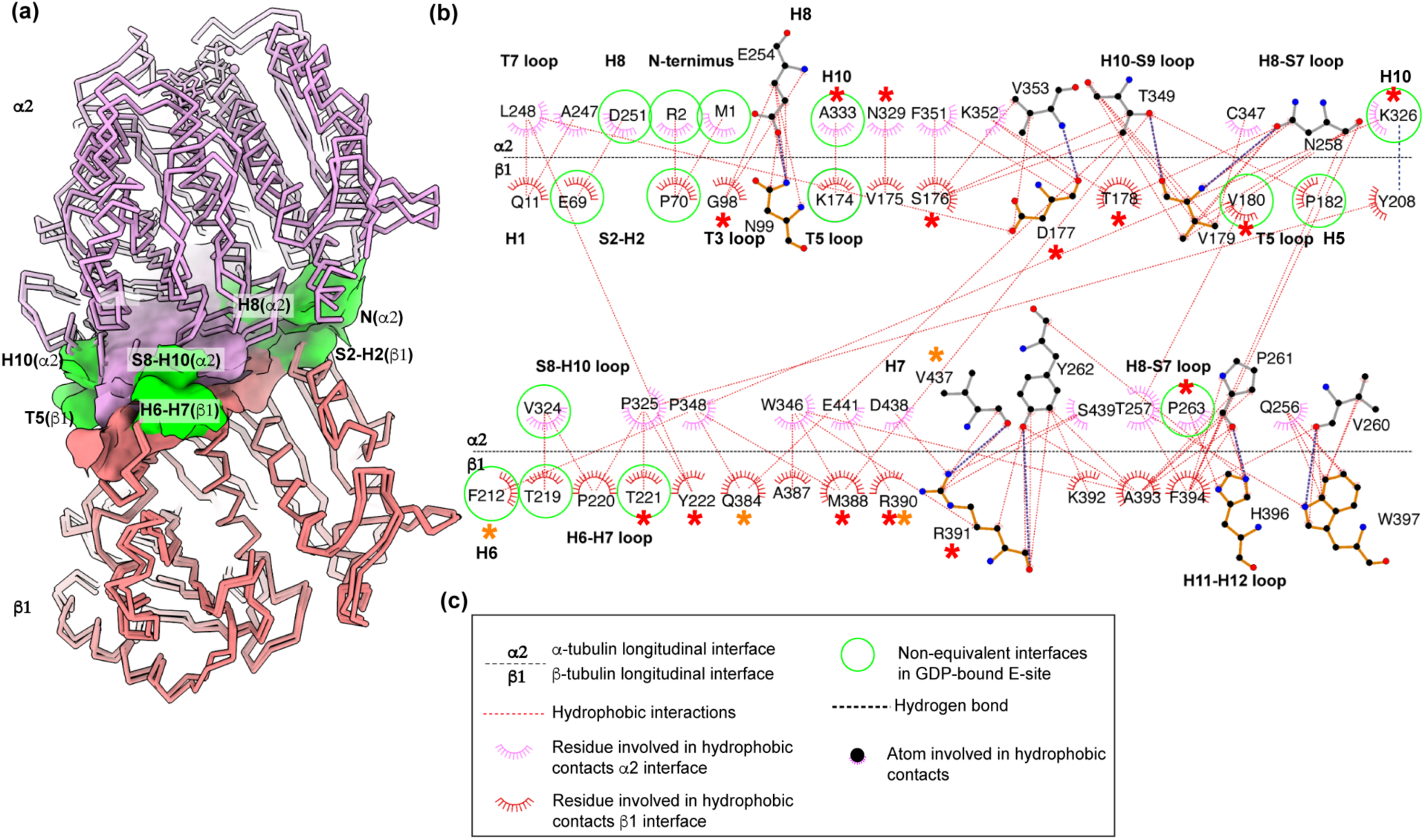
The longitudinal interface increases in the GDP-bound state compared to the GMPCPP-bound state. **(a)**. The α2 and β1-tubulin protomers from GMPCPP- and GDP-bound structures are shown as chain trace with the α2-tubulin, lilac and β1-tubulin, salmon. The two models are aligned on the rigid structural elements that coordinate the nucleotide base bound to β1-tubulin. Residues that make longitudinal contacts are shown as solvent-excluded surfaces. Residues found at the longitudinal interface in both GMPCPP-and GDP-bound microtubules are shown in lilac for α2-tubulin, and salmon, for β1-tubulin. Residues that participate in novel interactions at the GDP-bound longitudinal interface are shown in green. **(b)**. DIMPLOT^45,46^ showing a 2-D representation of molecular interactions at the longitudinal interface in the GMPCPP and GDP-bound states. Contacts found only in GDP microtubules highlighted by green circles. Residues mutated in human disease are highlighted by red stars^44^. Residues which are variable among tubulin isoforms are highlighted by orange stars (Fig. S3 and S4).

### The lateral interface is modulated by isoform-specific sequence variations

The lateral microtubule interface is substantially smaller than the longitudinal interface and is formed by a limited number of small loops: the M loop on one protofilament and the H1-H1’, H1’-S2, H2-S3 loops on the neighboring protofilament (Fig.5). The lateral interface is composed mostly of polar residues as seen in other microtubule structures ^3,4,6,11–13,17,30^ ^5^(Fig. 5). H283 in the M loop is the key residue at the lateral interface between α-tubulin protomers. It makes van der Waals interactions with K60 in H1’-S2 and H88 in H2-S3. H283 in H2-S3 also H-bonds with the backbone carbonyl of Q85 in the adjacent protofilament (Fig. 5(b)). In addition to this key contact, E284 and Q285 in the M loop H-bond with H88 in H2-S3 and the backbone carbonyl of E55 in H1’-S2, respectively (Fig. 5(b-c)), and K280 in the M-loop is within H-bonding distance to E90 in the H2-S3 loop (Fig. 5(b)). All these residues involved in α-tubulin lateral contacts are conserved among human tubulin isoforms. At the β-tubulin lateral interface Y281 in the M loop is the key residue, involved in van der Waals interactions with K58 in H1’-S2 and P87 in the H2-S3 loop, and a hydrogen bond with Q83 in the H2-S3 loop of the neighboring protofilament (Fig. 5(d)). The Y281 backbone carbonyl oxygen is also within H-bonding distance to R86 in the H2-S3 loop (Fig. 5(e)). Additional contacts are made between S278 in the M loop which H-bonds with D88 in the H2-S3 loop of the neighboring protofilament (Figs. 5(e)). Unlike the longitudinal interface, comparison of the lateral interface between the GMPCPP and GDP-bound structures did not show large conformational changes (Fig. 6(a-b)). Specifically, when we aligned the GMPCPP and GDP structures on the β1’-tubulin, the distances between residues at the lateral interface changed by less than 0.7 Å. This is in agreement with previous structures ^3,4^

**Figure 5.**
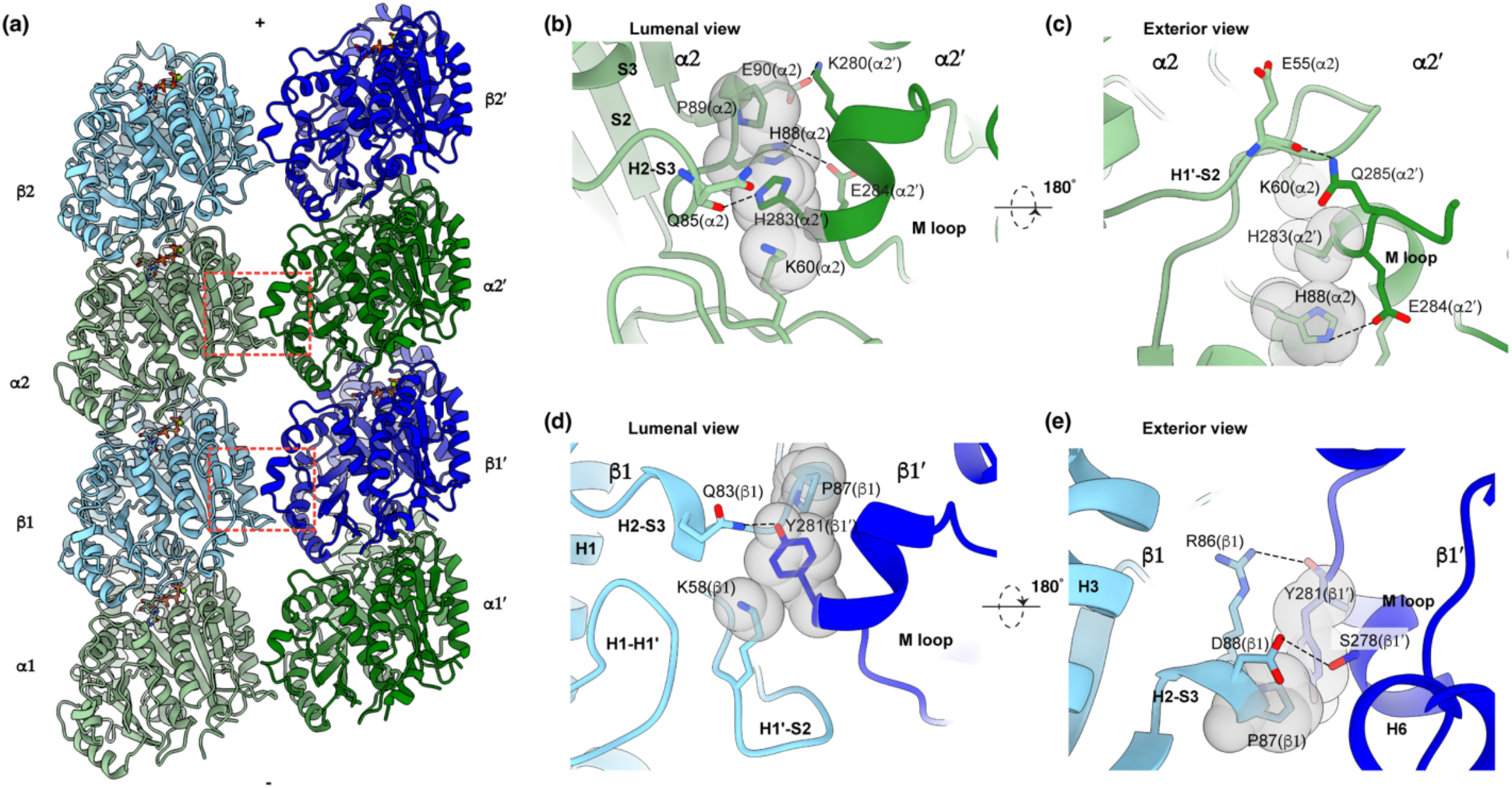
The α- and β-tubulin lateral interfaces. **(a)** Atomic model of two adjacent protofilaments in a GMPCPP-microtubule, view from the luminal side. β-tubulin, blue and α-tubulin, green. The lateral interfaces between α1-α1’ and β1-β1’ are highlighted by red dashed outlines. “+” and “-“ indicate the plus and minus ends, respectively. **(b-c)**. Interactions at the α-tubulin lateral interface; α1 and α1’ denote laterally adjacent α-tubulin protomers. **(d-e)**. Interactions at the β-tubulin lateral interface; β1 and β1’ denote laterally adjacent β-tubulin protomers. Hydrogen bonds shown as black dashed lines. Residues mutated in human disease highlighted with red stars^44^.

**Figure 6.**
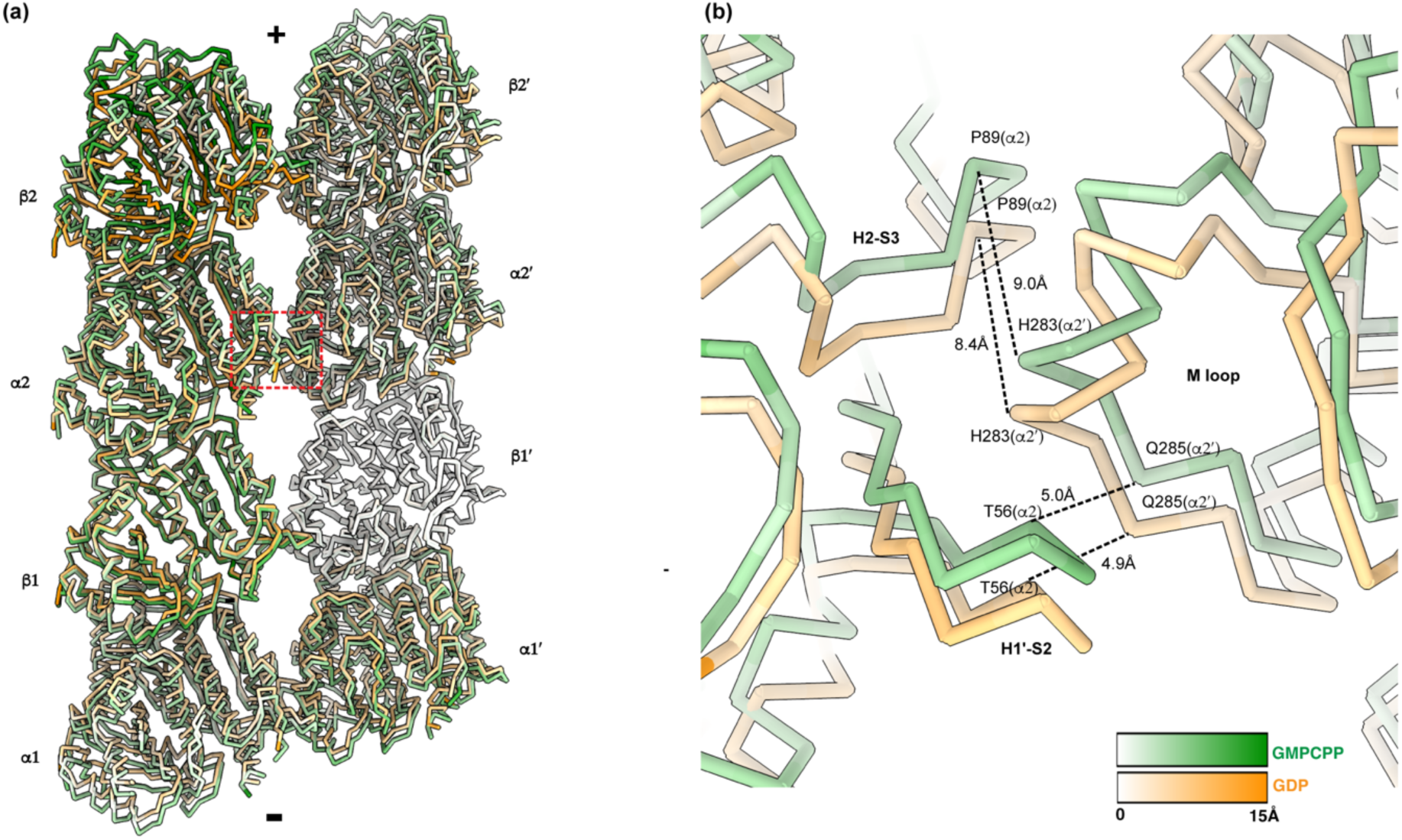
The lateral interface in the GMPCPP and GDP-bound states. **(a).** Two lateral protofilaments are shown as chain traces. The protofilaments were aligned on the β-sheet of NBD in β1’. Tubulin dimers are labeled as α1 and β1 at the minus end, “-“, and α2 and β2 at the plus end, “+“, in one protofilament; α1’ and β1’ at the minus end and as α2’ and β2’ at the plus end in the second protofilament. The lateral interface between α1 and α1’ is outlined with the red dashed line. Models are colored by RMSD between αCs with the GMPCPP structure colored on a gradient from white to bright green to dark green and the GDP structure colored on a gradient from white to yellow to orange. **(b)**. The lateral interface between α1 and α1’ does not change significantly between the GMPCPP and GDP-bound states. Distances between αC residues in the lateral loops are shown with black dashed lines.

While the longitudinal and α-tubulin lateral interfaces are stringently conserved among mammalian isoforms, the β-tubulin lateral interface concentrates much of the sequence variability (Fig. S4 and Fig. 7(a-b)), specifically in the M, H2-S3 and H1’-S2 loops, directly involved in lateral contacts, as well as structural elements which help position these loops at the lateral interface. Q83 in the H2-S3 loop, which H-bonds with Y281 in the M loop of the laterally adjacent β-tubulin is substituted to histidine in βIII (TUBB3) and to alanine in βVI (TUBB1). Mutation to alanine abrogates the H-bond with Y281, weakening this interface. This interface is further weakened in βVI (TUBB1) by the substitution of R86 to a smaller glutamine (Fig. 6(b), Fig. S4).

**Figure 7.**
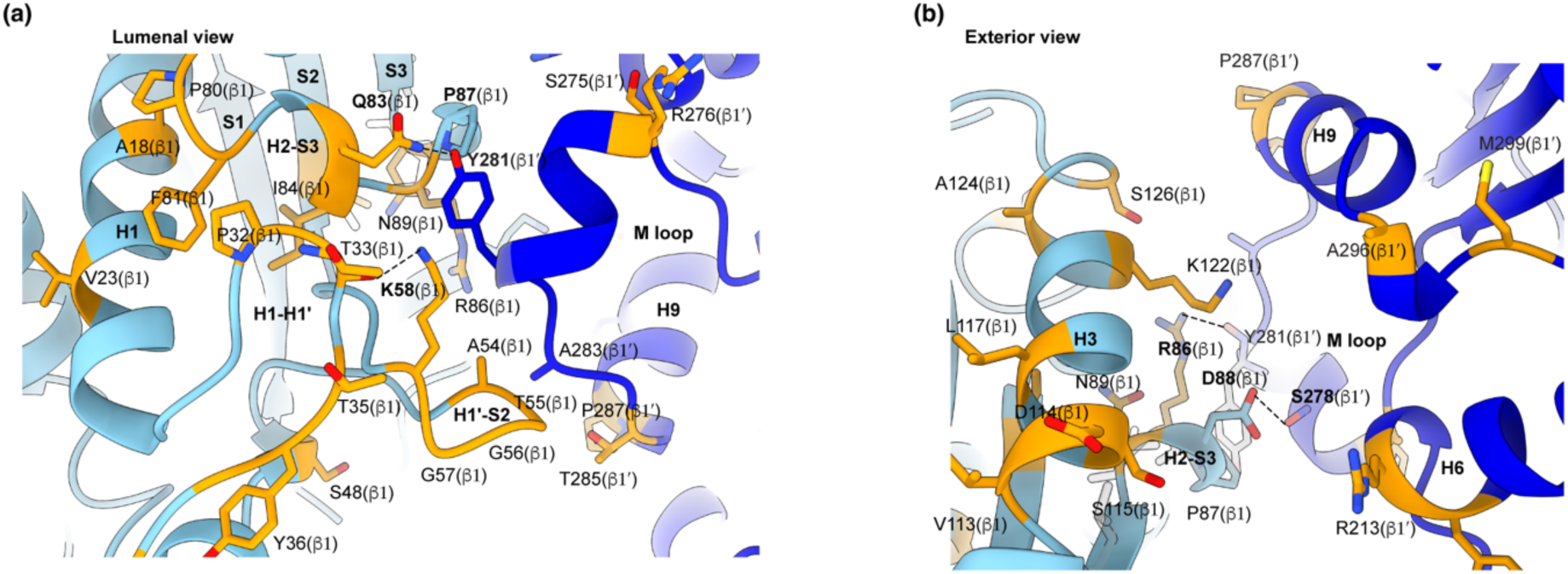
The lateral interface is modulated by isoform-specific sequence variations. **(a-b)**. Isoform variable residues at the β-tubulin lateral interface and concentration of sequence variability in the second shell of residues away from the β-tubulin interface. Residues directly involved in lateral contacts shown in bold. Residues that are variable among human tubulin isoforms shown in orange.

Perhaps because the penalty in fitness would be too high, the majority of tubulin isoform sequence variability occurs in residues that are not part of the lateral interface, but that instead help position the loops that form this interface (Fig. 7(a-b)). Replacement of T55 in the H1’-S2 loop to a tyrosine in βVI (TUBB1) can create additional contacts at the lateral interface with K362 in the S9-S10 loop and L284 in the M loop. G57 in the H1-S2 loop is replaced with bulkier residues in all isoforms other than βI(TUBB), βIVa/b(TUB4A/B) and βVIII(TUBB8) (Fig. S4, Fig. 7(a)), changing the flexibility of this loop and indirectly affecting the conformation of the M loop at the lateral interface. P80 is replaced by an alanine or a lysine in βIII(TUBB3) and βVI(TUBB1), respectively. This increase in backbone flexibility would impact the conformation of the H2-S3 loop and thus the positioning of Q83 at this interface (Fig. 7a). P32 and F81 pack against each other forming a rigid interface. In βVI (TUBB1) both are replaced by leucines (Fig. S4, Fig. 7(a)) which would increase the conformational flexibility and affect the positioning of the H1’-S2 and H2-S3 loops. Thus, isoform-specific substitutions modulate the flexibility and chemistry of lateral interface loops, and are likely responsible for the differences in dynamic properties of microtubules assembled from different human β-tubulin isoforms ^11–14^.

## Discussion

We report 2.9Å resolution structures of human α1B/βI+βIVb microtubules in GMPCPP- and GDP-bound states. Our reconstructions revealed that the N-terminal region of the α-tails interact in trans with the β-subunit of the neighboring tubulin dimer. These interactions reinforce the longitudinal interface, but the flexible nature of the tails could also enable tubulin subunits to stay tethered when the canonical interface is not fully formed or broken. Molecular dynamics revealed that that α-tail interacts with the canonical longitudinal interface, occluding it ^31^. Consistent with this, loss of the α-tubulin tail increases microtubule growth rates ^31^. Thus, the α-tail undergoes a switch from occluding the longitudinal interface in the tubulin dimer and inhibiting polymerization, to helping tether neighboring tubulin dimers to one another. While our structure visualized only a few of the residues at the N-terminus of the α-tail engaged in interactions with the β-tubulin body of the neighboring dimer, it is possible that the remaining negatively-charged α-tail forms transient, flexible interactions with the β-tubulin surface beyond what we visualized using cryo-EM, a technique not suited for capturing transient and conformationally variable interactions.

Our structures in GMPCPP- and GDP states also showed that α1B/βI+βIVb microtubules undergo similar remodeling and reinforcement of the longitudinal interface as microtubules assembled from porcine brain tubulin ^3–5^ or the human neuronal α1AβIII isoform ^6^, suggesting that this a property shared among mammalian tubulins. Comparative analysis of isoforms identified a concentration of sequence variability in the second shell of residues away from the β-tubulin lateral interface which can modulate the strength of this interface. Future high-resolution structures (< 3Å) of other tubulin isoforms will be needed for comparative studies aimed at understanding their different energetics of polymerization and depolymerization.

## Acknowledgements

We thank H. Wang (Multi-Institute cryo-EM Facility at the National Institutes of Health) for assistance with cryo-EM data collection. Image processing was performed on the Biowulf cluster maintained by the High Performing Computation group at the National Institutes of Health (NIH). A. R.-M. is supported by the intramural program of the National Institute of Neurological Disorders and Stroke and the National Heart, Lung and Blood Institute.

## Author Contributions

E.A.Z. collected and processed cryo-EM data. E.A.Z. with help from A.R.M. built models and analyzed structures. E.A.Z. and A.R.M. wrote manuscript. A.R-M. initiated, coordinated and supervised project.

## Accession Numbers

The GMPCPP-microtubule and GDP-microtubule models have been deposited at the Protein Data Bank under accession numbers PDB ID 8V2I and PDB ID 8V2J, respectively. Maps were deposited at Electron Microscopy Data Bank with accession numbers EMDB-42915, EMDB-42916, respectively.

## Methods

### Cryo-EM sample preparation

Tubulin from a human embryonic kidney (tsA201) cell line was purified as described previously^12,21^. GMPCPP bound microtubules were prepared as follows. A frozen aliquot of unmodified tubulin was thawed, and aggregates were removed by ultracentrifugation at 346,000 x g for 10 min at 4°C. Tubulin was diluted to 2 mg/mL in 1X BRB80 buffer (80mM PIPES, 1mM MgCl_2_, 1mM EGTA) and incubated in the presence of 1mM GMPCPP (Jena Bioscience) for 5 min on ice, and then polymerized at 37° C for 1 hour. Microtubules were pelleted by centrifugation at 126,000 x g for 10 min at 30° C. Supernatant was discarded and the microtubule pellet was washed with warm 1X BRB80 buffer. The microtubule pellet was re-suspended in cold 1X BRB80 buffer and placed on ice for 30 min to depolymerize. GMPCPP was added to 1 mM to the reaction while on ice for 5 min and then reaction was transferred to 37° C overnight for polymerization. The microtubule pellet was spun again at 126,000 x g for 10 min at 30° C, washed and resuspended in warm 1X BRB80. The concentration of polymerized microtubules was measured by denaturing in 6M Guanidine Hydrochloride and measuring absorbance at 280 nm. To prepare dynamic, GDP-bound microtubules, unmodified tubulin was precleared by centrifugation from aggregates as described above and then polymerized at 40μM in 1X BRB80 buffer and 2mM GTP for 30 min at 37° C for 30 min.

### Cryo-EM sample preparation and data collection

Double-cycled GMPCPP microtubules were diluted to 2.5μM in 1XBRB80 buffer, dynamic microtubules were left undiluted at 40 μM. 5 µls were applied to a glow discharged 1.2/1.3 CF-1.2/1.3-3Au-50 (Electron Microscopy Sciences, part # AUFT313-50) cryo-EM grid and allowed to absorb for 30 sec. The sample was blotted with Blotter Whatman® No.1 filter paper (Leica microsystems, No: 16706440) for 4 s using a Leica EM GP2 (Leica microsystems) with the chamber set to 30°C and 90% humidity and subsequently plunged into liquid ethane.

Data collection for GMPCPP-microtubules was performed on a Krios microscope (FEI) equipped with a K2 camera (Gatan) and an energy filter of 20 eV slit width. A total of 6,930 movies were acquired at the magnification of 130,000× in counting mode, physical pixel size 1.06 Å/pix, with a 70 μm C2 aperture, 100 μm objective aperture and spot size of 7. The dose rate was 8 e^-^/pix/sec and the total exposure time 7.8 sec for a 40-frame movie with a total electron dose of 56 e^-^/Å^2^. Data were collected in SerialEM (https://bio3d.colorado.edu/SerialEM/) with defocus values from -0.5 μm to -2.5 μm. Data collection parameters are shown in Table S1.

Data collection for GDP-microtubules was performed on a Krios microscope (FEI) equipped with a K3 camera (Gatan) and an energy filter of 20 eV slit width. A total of 8,856 movies were acquired at the magnification of 105,000× in super-resolution mode, physical pixel size 0.83 Å/pix, with a 70 μm C2 aperture, 100 μm objective aperture and spot size of 5. The dose rate was 20.22 e^-^/pix/sec and the total exposure time 1.58 sec for a 21-frame movie with a total electron dose of 46 e/Å^2^. Data were collected in SerialEM (https://bio3d.colorado.edu/SerialEM/) with defocus values from -0.6 μm to -2.4 μm. Data collection parameters are shown in Table S1.

### Electron microscopy image processing

All image processing was carried out in RELION v3.0^32^. Frames were aligned and summed using MotionCor2^33^ (Fig. S1, Table S1). The contrast transfer function (CTF) was estimated using GCTF^34^. Movie sums with Thon rings beyond 5 Å were selected for further image processing. Microtubules were manually selected. 124,408 overlapping segments with a box size of 420 pixels (GMPCPP data) or 167,963 overlapping segments with a box size of 480 pixels (GDP data) were extracted every 82 Å and down sampled to a pixel size 4.24 Å/pixel (GMPCPP data) or 4.98 Å/pixel (GDP data). Three-dimensional (3D) reconstruction of the microtubule was carried out using the previously described MiRP procedure (Fig. S1(a))^35^. Briefly, segment averages were generated for each microtubule and were subjected to supervised classification against 11 to 16 protofilament (PF) synthetic microtubule references, low pass filtered to 15 Å (Fig. S1(b)). 14 PF microtubule segments were chosen for further processing. Rotation angles and translations were reset to zero, Psi and Tilt angles were set to priors, followed by one round of refinement against the 14 PF synthetic reference. For each microtubule the most common Rotation angle was derived and assigned to all segments in that microtubule, followed by 3D refinement against the 14 PF synthetic reference with local sampling of Euler angles. Next, X/Y shifts were smoothed to remove mistranslations along microtubules, followed by refinement. Particles were re-centered and re-extracted and segment averages were generated for each microtubule. Seam location along each microtubule was assigned by performing a supervised 3D classification without alignment against 28 synthetic references with all possible seam positions. Each microtubule was then assigned to a common class, followed by adjustment of angles and translations along the helical axis. Unbinned microtubule segments were re-extracted and re-centered. 3D refinement with and without imposing symmetry was performed. The quality of the resulting map was further improved by performing protofilament refinement^23^. Briefly, the signal was subtracted from all but a single protofilament in each microtubule segment. The process was repeated for each protofilament, resulting in 14x times more particles. The protofilament particles were refined as single particles. The nominal resolution of the maps is 2.9 Å (FSC cutoff 0.143) (Fig. S1 (d-e). Lattice distortions were quantified from the refined rotation angle values between adjacent protofilaments (Figs. S2(a-b)). The maps were sharpened in RELION^32^ with B-factor -75 Å^2^ (GMPCPP-MT) or -62 Å^2^ (GDP-MT) or modified with DeepEMhancer^36^. Local resolution was estimated using Monores (Fig. S1(e))^37^. Structures were analyzed in UCSF Chimera^38^ and UCSF ChimeraX^39^. Figures were made in UCSF ChimeraX^39^.

### Model building and refinement

The cryo-EM-derived structure of a human unmodified microtubule (PDB ID: 5N5N)^12^ was used as an initial model to build the GMPCPP and GDP-bound microtubule structures. The initial model was rigidly docked into the cryo-EM map using the UCSF Chimera “Fit in Map” tool, followed, by iterative rounds of refinement in Phenix^40^ and manual rebuilding in Coot^41^, software packages maintained within the SBGrid platform ^42^. The iterative refinement was performed against two unsharpened half maps to avoid overfitting. The quality of the final models were validated using the Phenix Comprehensive Validation tool (Table S2).

### Dimer repeat distance analysis

To calculate the dimer repeat distances, each of the two cryo-EM datasets were split into three subsets and used for three independent reconstruction using the MiRP procedure^35^. For each reconstruction, the helical symmetry parameters, rise and rotation per subunit, were refined using the RELION helical reconstruction procedure^43^. The refined rise per subunit was multiplied by 14 (protofilament number in the reconstruction) and divided by 1.5 (3-start helix). The individual values are listed in Fig. S2(c)).

