## Supplementary information for "Cryo-EM structures of human α1B/βI+βIVb microtubules shed light on isoform specific assembly"

#### Supplementary tables.

#### Supplementary Table 1. Cryo-EM data collection and processing

| | | |
|---|---|---|
| **Data collection** | | |
| | **MT-GMPCPP data** | **MT-GDP data** |
| Microscope | Krios | Krios |
| Detector | K2 | K3 |
| Data collection software | Serial EM | Serial EM |
| Nominal magnification | 130,000x | 105,000x |
| Voltage (kV) | 300 | 300 |
| Electron exposure ($e^-$/Å$^2$) | 56 | 46 |
| Exposure rate ($e^-$/pixel/s) | 8.0 | 20.2 |
| Defocus range (µm) | -0.5 to -2.5 | -0.6 to -2.4 |
| Physical pixel size (Å) | 1.06 | 0.83 |
| Time per frame (s) | 0.195 | 0.075 |
| Total exposure time (s) | 7.8 | 1.58 |
| Frames per movie | 40 | 21 |
| Movies collected | 6,930 | 8,856 |
| **Image processing** | | |
| | **Microtubule-GMPCPP map** (EMDB-42915) | **Microtubule-GDP map** (EMDB-42916) |
| Symmetry imposed | C1 | C1 |
| Initial particle images | 124,408(microtubule segments) | 167,962(microtubule segments) |
| Final particle images | 1,293,027(protofilament particles) | 556,978(protofilament particles) |
| Map resolution (Å) | 2.9 | 2.9 |
| FSC threshold | 0.143 | 0.143 |
| Map resolution range (Å) | 2.5 to 3.5 | 2.5 to 3.5 |
| Map sharpening B factor (Å$^2$) | -75 | -62 |

**Supplementary Table 2. Model refinement and validation statistics**

| | **microtubule-GMPCPP model** (EMDB-42915) (PDB 8V2I) | **microtubule-GDP model** (EMDB-42916) (PDB 8V2J) |
|---|---|---|
| Atomic modeling refinement packages | Phenix, Coot | Phenix, Coot |
| Initial model used (PDB code) | 5N5N | 5N5N |
| Model resolution (Å) | 2.9 | 2.9 |
| FSC threshold | 0.143 | 0.143 |
| Model composition | | |
| Non-hydrogen atoms | 13,623 | 13,580 |
| Protein residues | 1,732 | 1,732 |
| Ligands | 2 GTP; 2 GMPCPP; 4 $Mg^{2+}$ | 2 GTP; 2 GDP, 2 $Mg^{2+}$ |
| *B* factors ($Å^2$) | | |
| Protein | 122 | 118 |
| Ligand | 107 | 106 |
| Bonds (RMSD) | | |
| Bond lengths (Å) | 0.006 | 0.006 |
| Bond angles (°) | 1.240 | 1.231 |
| Validation | | |
| MolProbity score | 1.13 ($99^{th}$ percentile) | 1.21 ($99^{th}$ percentile) |
| Clashscore | 2.25 ($99^{th}$ percentile) | 2.18 ($99^{th}$ percentile) |
| Poor rotamers | 4 (0.28%) | 3 (0.21%) |
| EMRinger score | 3.13 | 2.80 |
| Ramachandran plot | | |
| Favored | 1,686 (97.33%) | 1,673 (96.57%) |
| Allowed | 46 (2.67%) | 59 (3.43%) |
| Disallowed | 0 | 0 |
| Rama-Z score | | |
| whole | 0.33 | 0.18 |
| helix | 1.22 | 1.09 |
| sheet | 0.76 | -0.74 |
| loop | -1.06 | -0.54 |
| Model vs. Data | | |
| CC(volume) | 0.85 | 0.88 |
| CC(mask) | 0.85 | 0.87 |

### Supplementary Figures.

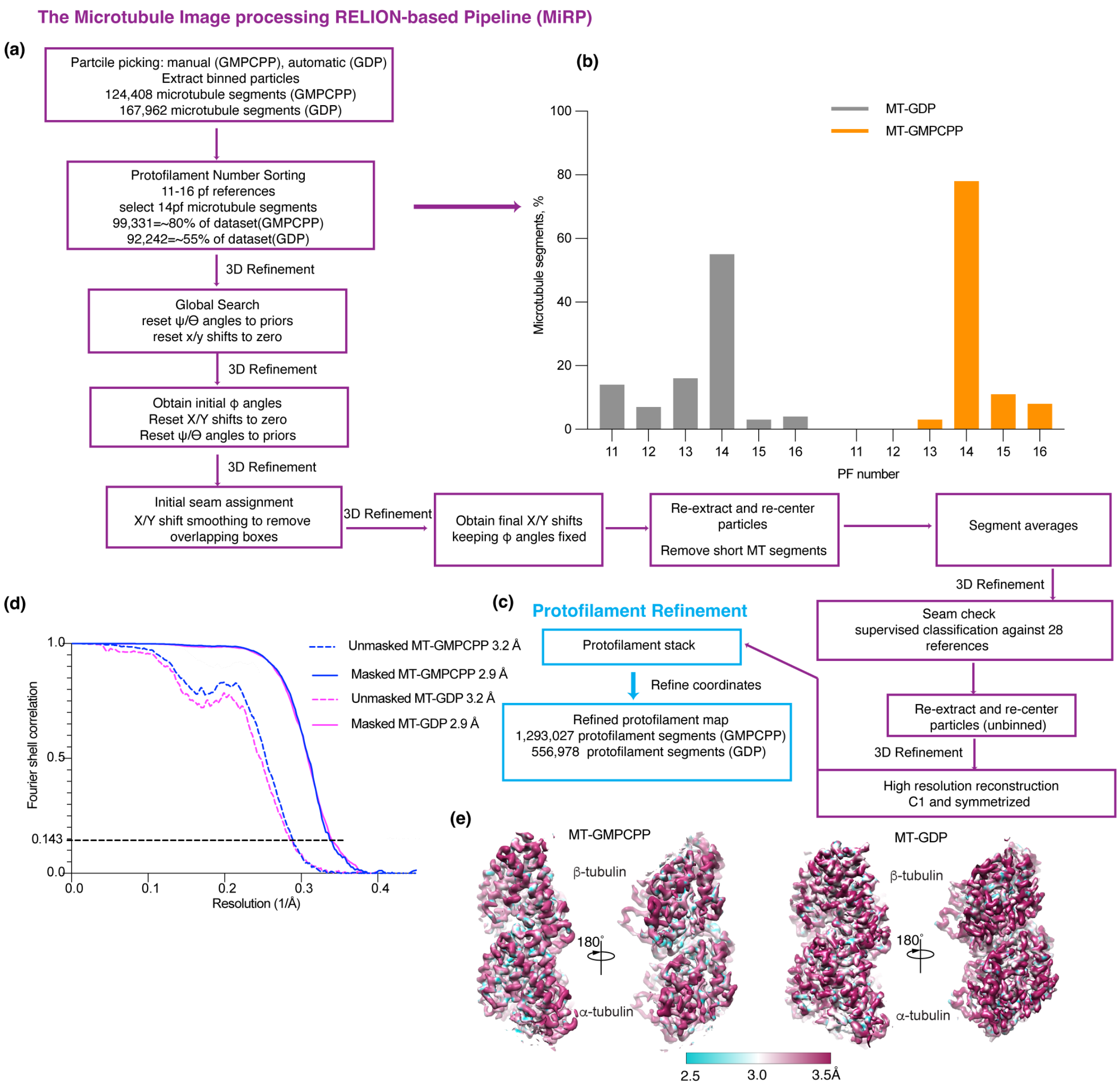


**Supplementary Figure 1. Cryo-EM image processing.**

**(a)**. Initial image processing was performed following the MiRP protocol[1] (Methods). **(b)**. Protofilament number distribution for GMPCPP and GDP-bound microtubules. **(c)**. The protofilament refinement procedure [2] was used to obtain the final high-resolution maps for 14-protofilament microtubules (Methods). **(d)** Fourier shell correlation (FSC) curves for each microtubule reconstruction, masked and unmasked, show their nominal resolution. FSC estimates were calculated at 0.143 criterion. **(e)**. Local resolution estimates show the resolution in the tubulin core reaches 2.5Å.

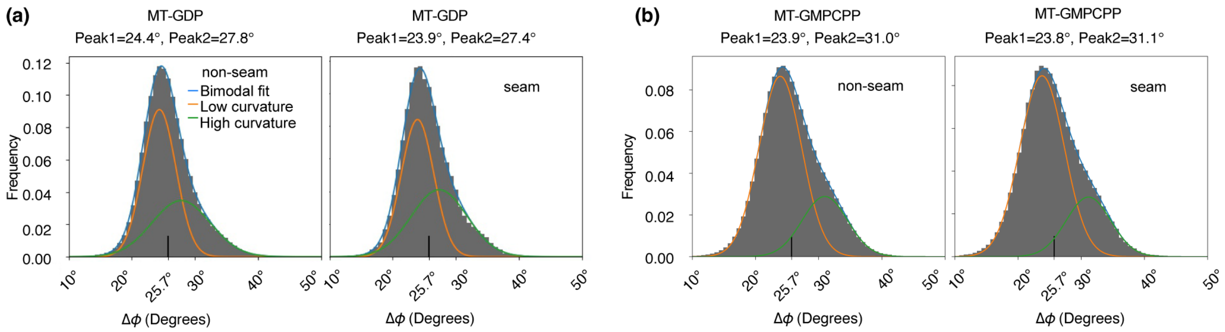


(c)

| | MT-GDP | MT-GMPCPP |
|---|---|---|
| **Dimer repeat distance 1** | 81.67 Å | 84.31 Å |
| **Dimer repeat distance 2** | 81.67 Å | 84.15 Å |
| **Dimer repeat distance 3** | 81.67 Å | 84.19 Å |
| **Mean±SD** | 81.67±0.00 Å | 84.22±0.08 Å |

**Supplementary Figure 2. Distribution of rotation angles and axial dimer repeat values in GMPCPP and GDP-bound microtubules.**

**(a).** Histogram of rotation angles (φ angle) between adjacent protofilaments for GDP-bound microtubule shows two peaks, a major peak of 24.4º and a minor peak of 27.8º for non-seam protofilaments, and a major peak of 23.9º and a minor peak of 27.4º for protofilaments at the seam. **(b).** Histogram of rotation angles (φ angle) between adjacent protofilaments for GMPCPP-stabilized microtubule shows two peaks, a major peak of 23.9º and a minor peak of 31.0º for non-seam protofilaments, and a major peak of 23.8º and a minor peak of 31.1º for protofilaments at the seam. These angles deviate from a lattice with symmetric protofilament geometry which would have an angle of 25.7° ($\varphi_{sym}$=360°/N, N is the number of protofilaments). **(c)**. Dimer repeat distance for GMPCPP- and GDP-bound microtubules, each obtained from independent reconstructions using 1/3 of the data (Methods).

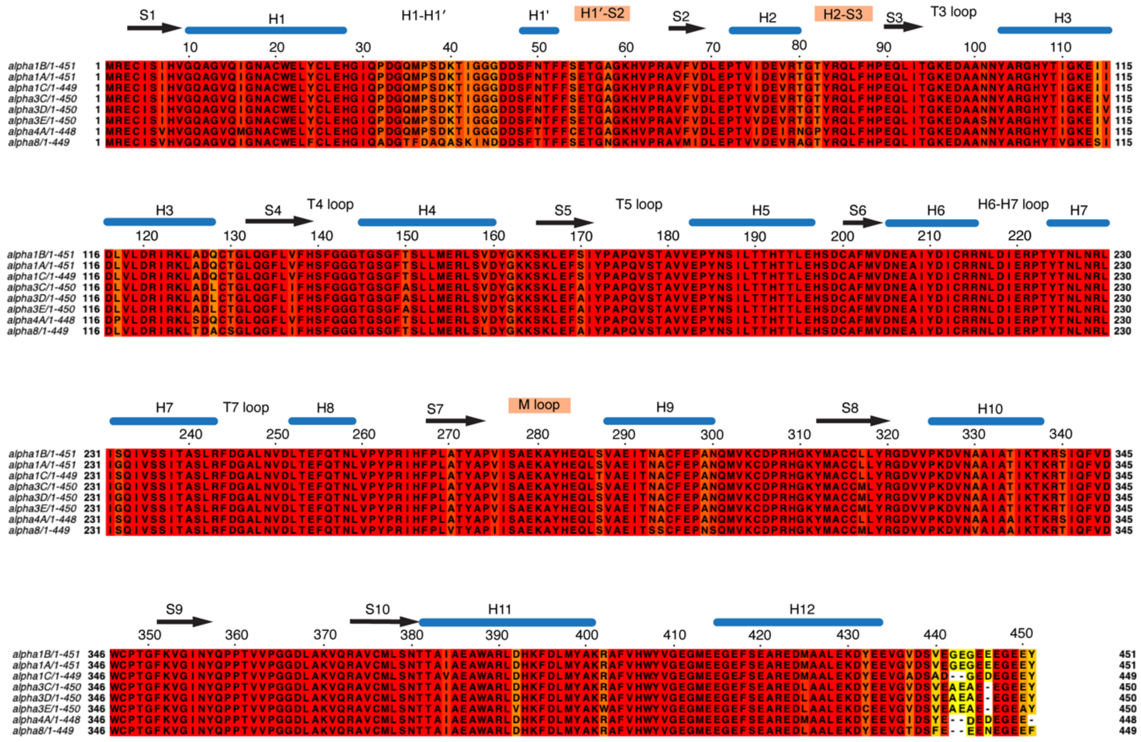

**Supplementary Figure 3. Sequence alignment of human α-tubulin isotypes.**

Sequence numbering corresponds to αIB-tubulin (TUBA1B) sequence UniProt P68363. β-strands, black arrows; helices, blue lines; lateral loops are highlighted by salmon rectangles. Sequences were aligned using Jalview Muscle with default settings[3].

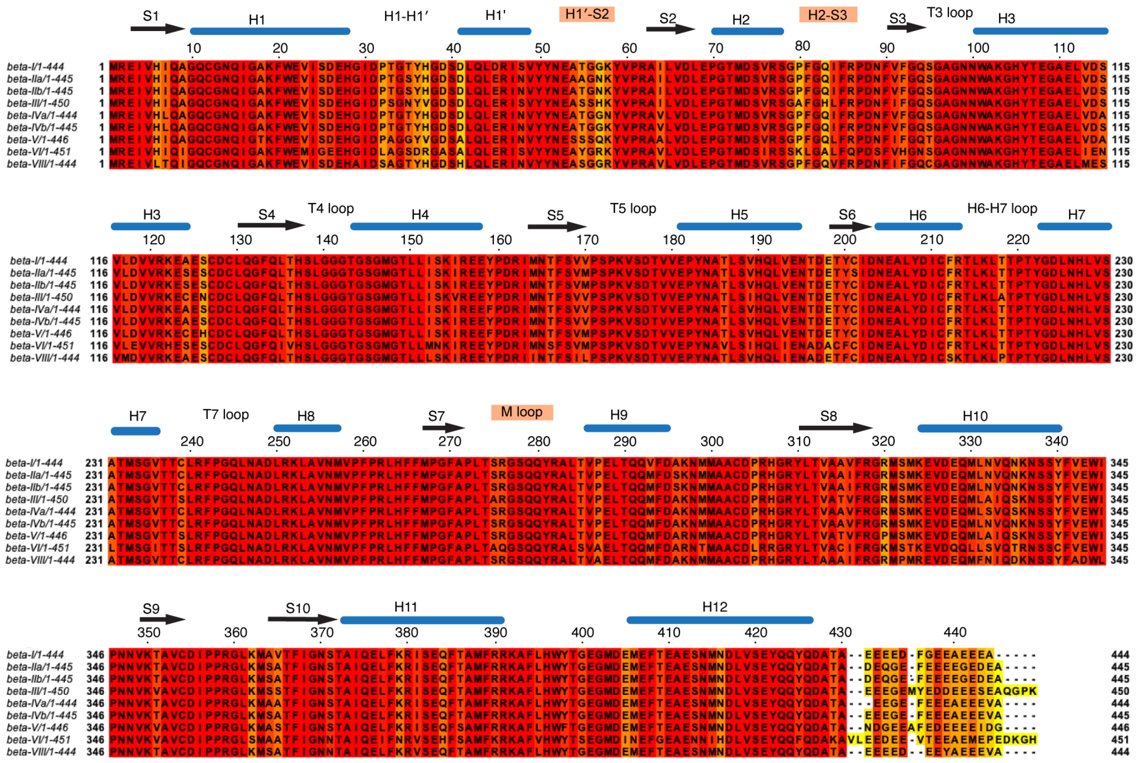

**Supplementary Figure 4. Sequence alignment of human β-tubulin isotypes.**

Sequence numbering corresponds to βI-tubulin (TUBB) sequence UniProt P07437. β-strands, black arrows; helices, blue lines; lateral loops are highlighted by salmon rectangles. Sequences were aligned using Jalview Muscle with default settings[3].

### References.


1. Cook, A.D., Manka, S.W., Wang, S., Moores, C.A., and Atherton, J. (2020). A microtubule RELION-based pipeline for cryo-EM image processing. J Struct Biol *209*, 107402. 10.1016/j.jsb.2019.10.004.
2. Debs, G.E., Cha, M., Liu, X., Huehn, A.R., and Sindelar, C.V. (2020). Dynamic and asymmetric fluctuations in the microtubule wall captured by high-resolution cryoelectron microscopy. Proc Natl Acad Sci U S A *117*, 16976-16984. 10.1073/pnas.2001546117.
3. Waterhouse, A.M., Procter, J.B., Martin, D.M., Clamp, M., and Barton, G.J. (2009). Jalview Version 2--a multiple sequence alignment editor and analysis workbench. Bioinformatics *25*, 1189-1191. 10.1093/bioinformatics/btp033.
